## Supplementary material for "Integrated information structure collapses with anesthetic loss of conscious arousal in *Drosophila melanogaster*": Text S1

**Text S1. Effect of anesthesia is consistent at different binarization thresholds**

In the main text, we computed system-level integrated information and the integrated information structure (IIS) after first binarizing LFPs at each epoch based on the median voltage. While we can potentially operationalise the states of the brain signals in many different ways, binarization at the median is the simplest discretisation process, and also normalizes entropy across all epochs. This is important as it controls for potential changes in entropy levels between wakefulness and anesthesia [1].

To exclude the possibility that our results vary wildly depending on the specific threshold used for binarization, we used different binarization thresholds and computed system-level integrated information for sets of 2 channels at a time. We found the effect of anesthesia to be consistent across thresholds for all flies (Fig S1).

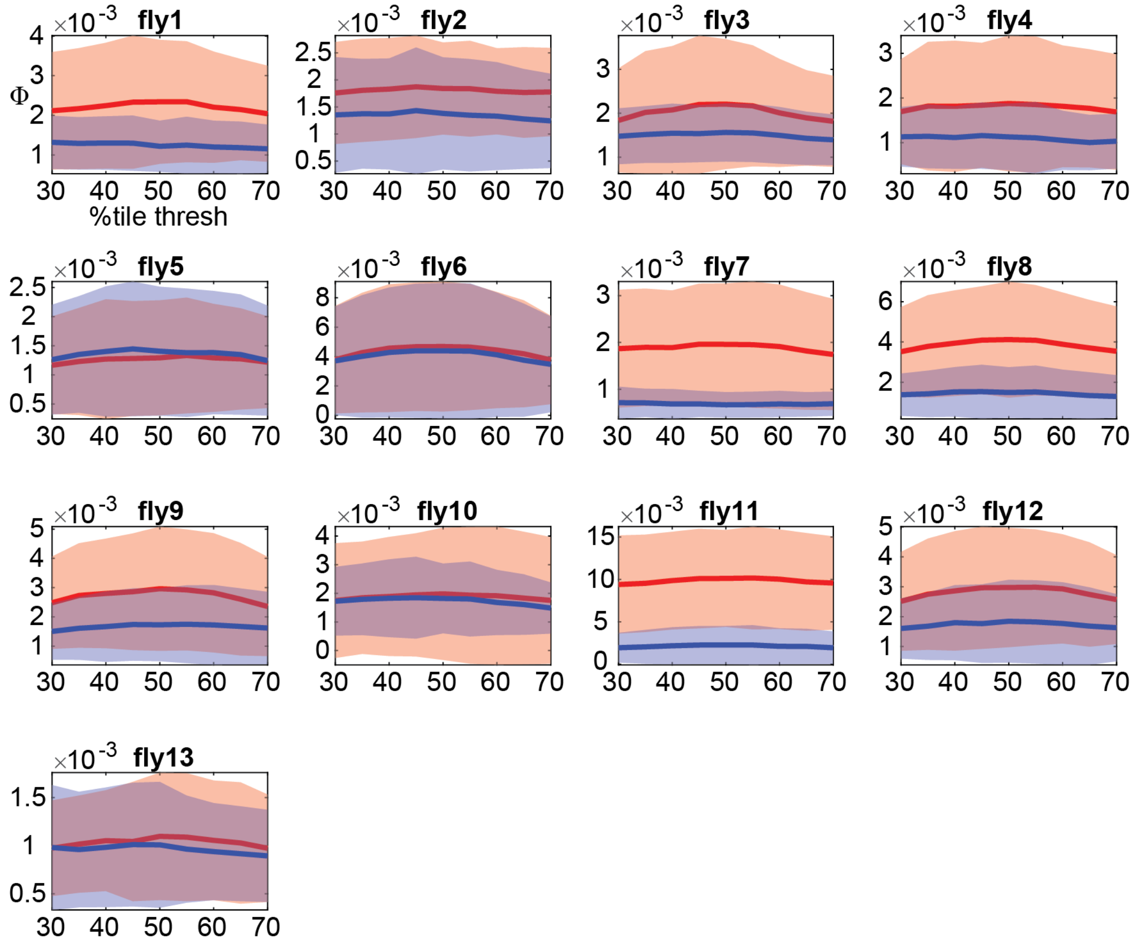


**Fig S1.** Effect of anesthesia on system-level integrated information is consistent across different binarization thresholds. We computed system-level integrated information (for sets of 2 channels at a time) after binarizing voltages of each channel at a given threshold (30th up to 70th percentiles in steps of 5; voltages become ‘1’ if above the threshold, and ‘0’ otherwise). Plotted is mean and standard deviation (across 105 channel sets per fly) for wakeful (red) and anesthesia (blue) conditions.
