## Supplementary material for "Integrated information structure collapses with anesthetic loss of conscious arousal in *Drosophila melanogaster*": Text S2

**Text S2. Effect of anesthesia is consistent using different timesteps**

In the main text, we computed system-level integrated information and integrated information structures (IIS) from TPMs built at the timescale $\tau$ = 4 ms. We chose this timescale based on the known physiology of synaptic interactions between neurons. Specifically, if $\tau$ is too small, it will not capture causal interactions that maximise integrated information [1–3]. Thus we chose 4 ms as our timescale. A comprehensive search across $\tau$ values is infeasible due to the computational cost of system-level integrated information.

To exclude the possibility that our results vary wildly depending on the specific $\tau$ value selected, we selected a random sample of 200 channel sets (out of the total 1365 channel sets), and recomputed system-level integrated information and the IIS at two other $\tau$ values, 2 ms and 6 ms. As can be seen in Fig S2 and Fig S3, the effect of anesthesia remains consistent with what we report in the main text (Fig 4 and Fig 5).

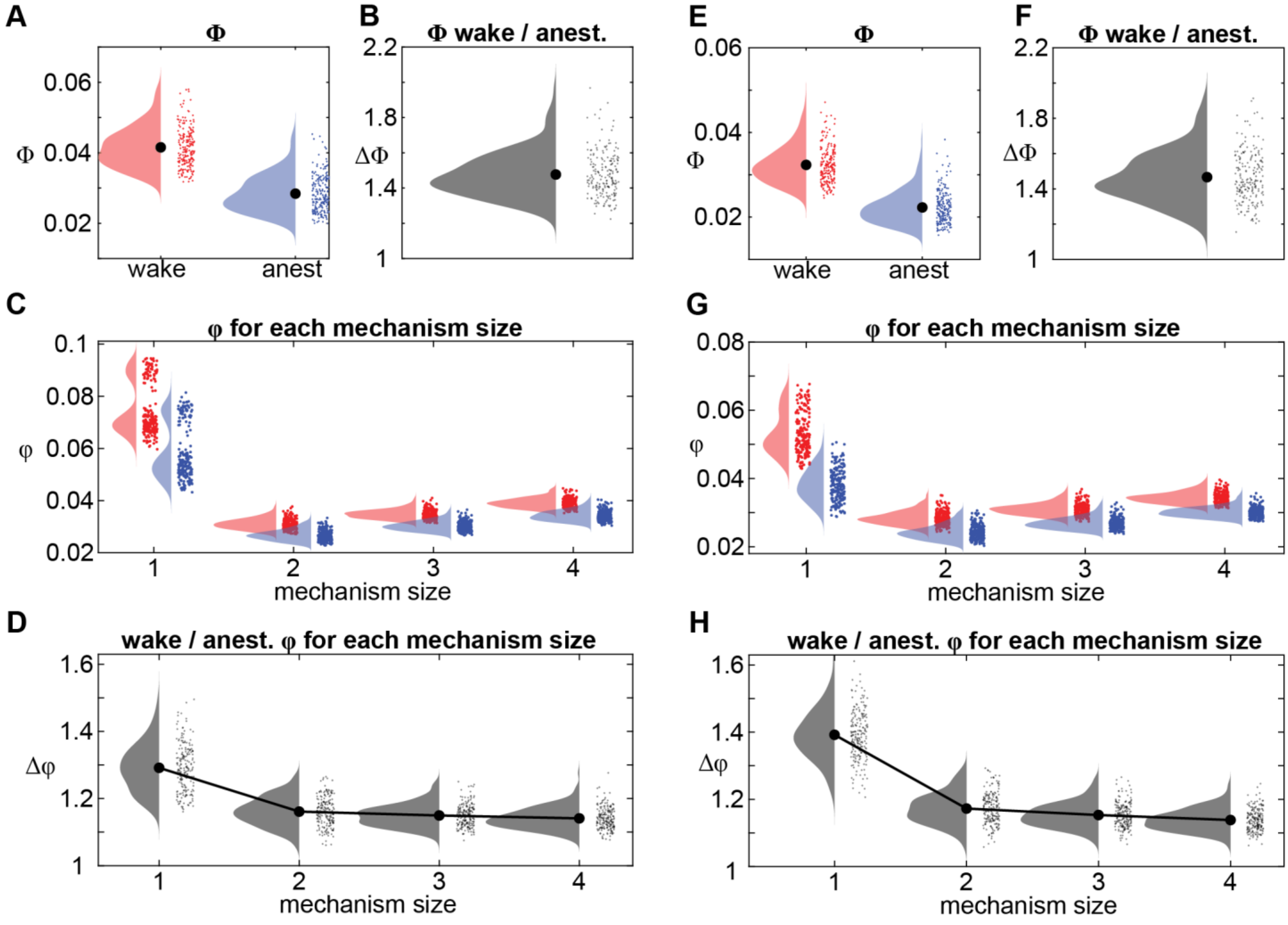

**Fig S2.** Effect of anesthesia on system-level integrated information (Φ) and the IIS is consistent for similar $\tau$ values. Format is the same as Fig 4 in the main article. (**A**-**D**) Results with $\tau$ = 2 ms. (**E**-**H**) Results with $\tau$ = 6 ms.

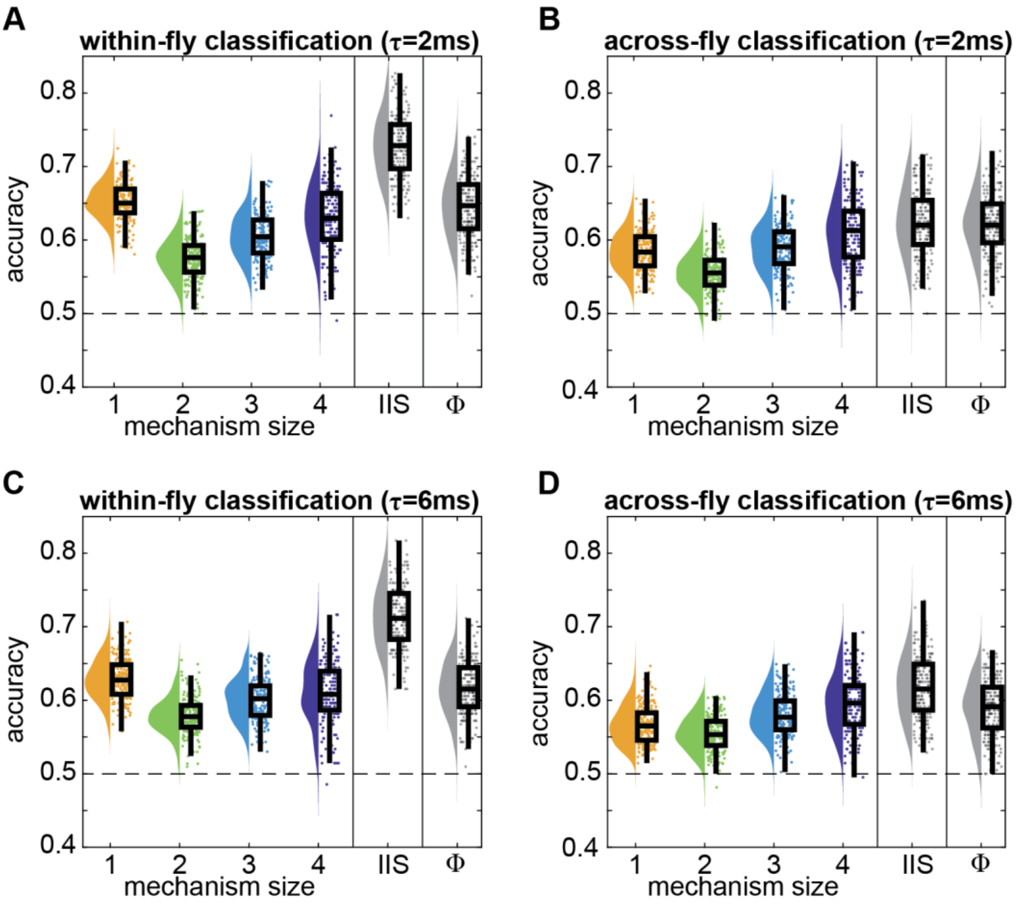


**Fig S3.** Classification accuracy of wake and anesthesia is consistent for similar $\tau$ values. Same format as Fig 5A-B in the main text. (**A**-**B**) Results with $\tau$ = 2 ms, (**C**-**D**) Results with $\tau$ = 6 ms.
