## Supplementary material for "Integrated information structure collapses with anesthetic loss of conscious arousal in *Drosophila melanogaster*": Text S3

**Text S3. “Disconnection” through statistical noising**

In the main text, we assessed the irreducibility of mechanisms (and of the system), by “disconnecting” connections between the mechanism and its purview such that some part of the mechanism affects only some part of the purview (and same for their complements). We carry out this “disconnection” by statistically noising the connections between the mechanism and the purview. Here we provide an example of this procedure in detail.

Consider a mechanism consisting of two channels, A and B, and a purview, consisting of channel C. For simplicity, we consider a case where both A and B have to be ‘1’ simultaneously to make C take the state ‘1’ at the next time step. This system’s state-by-channel (i.e., AB-by-C) TPM is shown in Fig S4A.

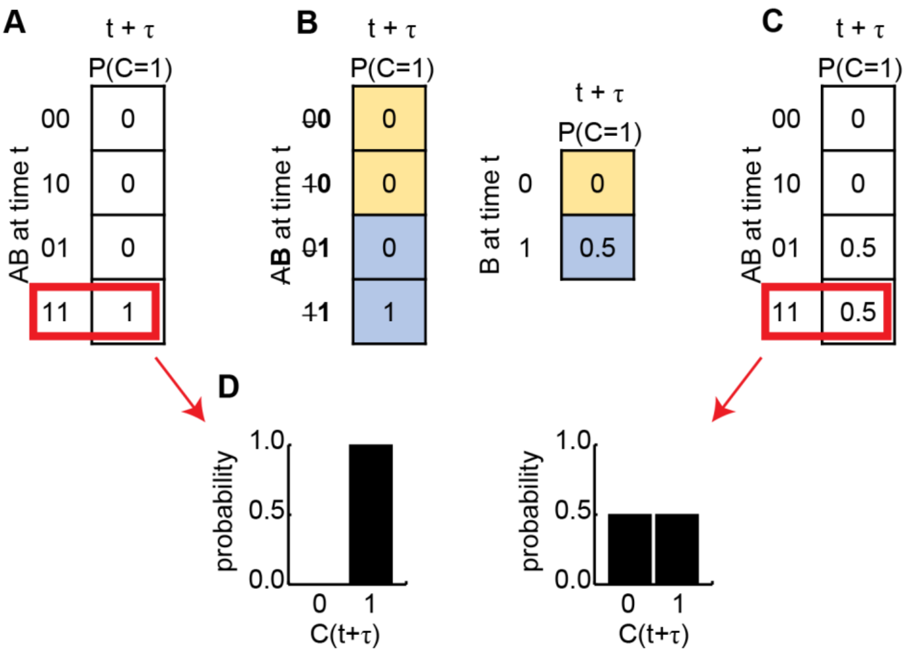


**Fig S4.** We estimate the effects of “disconnections” through statistical noising of the connection (not through physical disconnection). (**A**) An example state-by-channel transition probability matrix (TPM) for a mechanism AB and purview C. C becomes ‘1’ at time $t{+}\tau$ if both A and B are ‘1’ at time $t$. (**B**) To “disconnect” A from C, we replace the connection from A with noise by marginalising across the states of A. Colors indicate marginalising within each state of B. (**C**) Expanding the TPM marginalised over states of A returns the TPM to the original space of states of both A and B. (**D**) For the state of AB = 11 in red boxes in panel A and C, we obtain the probability distributions of C, before and after the imposed disconnection. We compare these distributions (using earth mover's distance) to obtain integrated information.

To assess the irreducible effect of AB on C, we carry out a "disconnection" (i.e. noising). Following the process as illustrated in Fig 1D-E, we want to compare the probability distribution of the purview, $P$(C=1) at $t+\tau$, when both A($t$) and B($t$) are known, to when only A($t$) is known or only B($t$) is known. To consider the case when only B($t$) is known, we replace A($t$) with noise, by marginalising over the possible states of A($t$). This gives us a disconnected TPM, as in Fig S4B. If we expand the disconnected TPM to again consider the possible states of B($t$) (which now give no information at all about C($t+\tau$), due to the prior marginalisation), we obtain Fig S4C.

Then, for a given state (e.g. AB='11'), IIT 3.0 uses earth mover's distance (EMD) to quantify the distance between the original probability distribution of C($t+\tau$), from the original TPM, and the probability distribution of C($t+\tau$) from the "disconnected" TPM where knowledge of some part of the mechanism has been factored out (in this example, knowledge of B($t$); Fig S4D).
