## Supplementary material for "Integrated information structure collapses with anesthetic loss of conscious arousal in *Drosophila melanogaster*": Text S4

### Text S4. Effect of anesthesia on system-level integrated information for each fly

In the main text, we compared the IIS to system-level integrated information, across all flies. Here, we show the effect of anesthesia on system-level integrated information per fly (Fig S5).

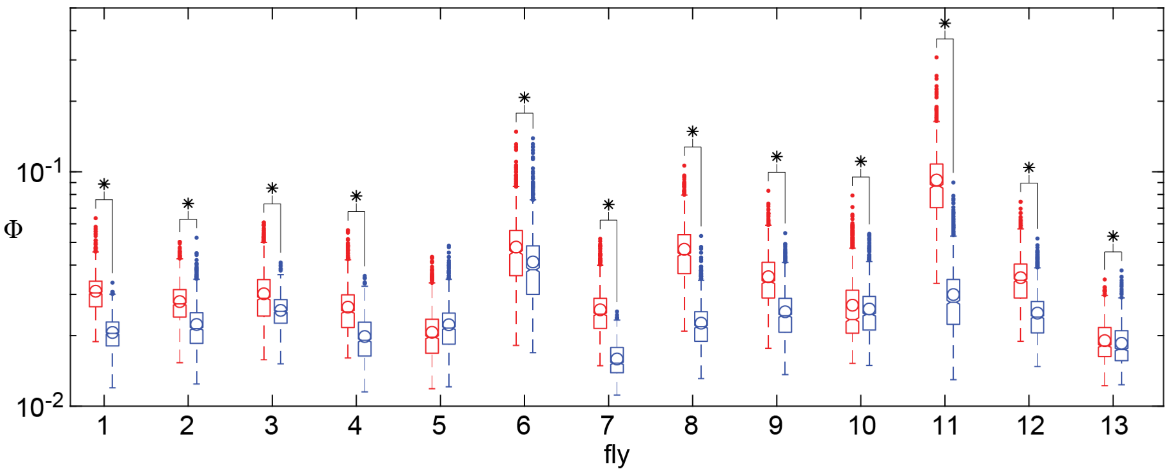


**Fig S5.** System-level integrated information, $\Phi$ (in log scale), during wakefulness (red) and anesthesia (blue) per fly. Boxes indicate 25th, 50th, and 75th percentiles across 1365 channel sets per fly. Circles indicate the mean across channel sets. Asterisks indicate significant one-tailed t-tests (system-level integrated information greater during wakefulness) across channel sets, $p < .001$.
