## Supplementary material for "Integrated information structure collapses with anesthetic loss of conscious arousal in *Drosophila melanogaster*": Text S5

**Text S5. IIS best predicts wakeful vs. anesthesia states**

In the main text, to assess the utility of the integrated information structure (IIS) in classifying wakeful vs. anesthetized states, we trained and tested support vector machines (SVMs) on either system-level integrated information (1 feature), the IIS (15 features), or integrated information of individual mechanisms (1 feature). While we found that SVMs trained on the IIS generally outperformed others, this might have been simply due to the IIS having more features to train on.

To exclude such a trivial interpretation, we conducted a complementary analysis with logistic regression, where we systematically compared the goodness of fits among models using an information theoretic model selection procedure (Akaike Information Criterion; AIC; [1]). AIC is defined as:

(S1) $\text{AIC} = -2ln(L) + 2k$

where $L$ is the maximum value of the likelihood function for the model and $k$ is the number of fitted parameters in the model. As the likelihood increases, the first term on the right-hand side becomes smaller, thus a smaller AIC is favoured. However, as a model includes more parameters, it gets penalised by the second term on the right-hand side, as the $2k$ term becomes larger. Thus, given two models with equal likelihoods, AIC selects the model with fewer parameters. Using AIC, we took into account the number of regressors (specifically, models with more regressors are penalised) and compared different model architectures (SII vs. other models with different numbers of mechanisms associated with integrated information).

We used the MATLAB implementation of logistic regression (fitglm.m) to regress a binary level of arousal (either wakeful or anesthetized) onto integrated information values. Specifically, we regressed the arousal level onto either 1) system-level integrated information (giving a single regressor, excluding the intercept), 2) the full IIS, where we used all 15 integrated information values associated with all mechanisms (giving 15 regressors) or 3) integrated information values of one of 1-, 2-, 3-, or 4-channel mechanisms (respectively giving 4, 6, 4, or 1 regressors). As a null model, we also regressed the arousal level onto only an intercept.

To interpret the results in the context of our SVM classification (Fig 5A in main text), we built a model per fly using all 1365 channel sets as observations (2 × 1365 observations per model; Fig. S6). The results were consistent with our conclusion with the SVM classification: the IIS performed better than the system-level integrated information even after accounting for the number of available variables fitted.

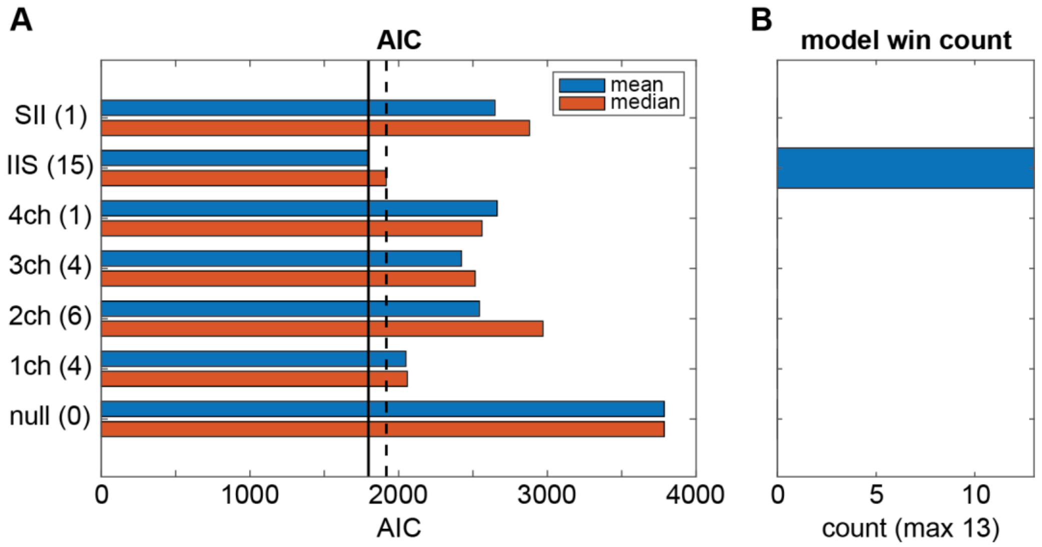


**Fig S6.** AIC values for logistic regression of the binary level of arousal onto different mechanism sizes, and system-level integrated information. Null models regressed the level of arousal onto only an intercept. (**A**) One AIC value was obtained from fitting models from 1365 channel sets per fly (observations per model = 1365 channel sets × 2 conditions). Shown are mean (blue) and median (red) of the 13 AIC values obtained from each of 13 flies. Solid and dashed lines indicate that the full IIS model performed the best (i.e. gave the smallest AIC) in terms of the mean of the median. Y-labels give the features used for fitting the model and the associated number of coefficients fitted (in parentheses), excluding the intercept. (**B**) The full IIS model was chosen as the best model, which gave the minimal AIC in all 13 flies, while the SII model was never chosen as the best model.
