## Supplementary material for "Integrated information structure collapses with anesthetic loss of conscious arousal in *Drosophila melanogaster*": Text S6

**Text S6. 1-channel mechanisms do not drive classification performance of the IIS**

As shown in Fig 4C, 1-channel mechanisms were associated with a higher magnitude of integrated information compared to other higher-order mechanisms. To quantify the degree of contribution of these 1-channel integrated information for the IIS classification, we repeated the classification analysis without integrated information associated with 1-channel mechanisms (Fig S7), which demonstrated no substantial contribution of 1-channel integrated information.

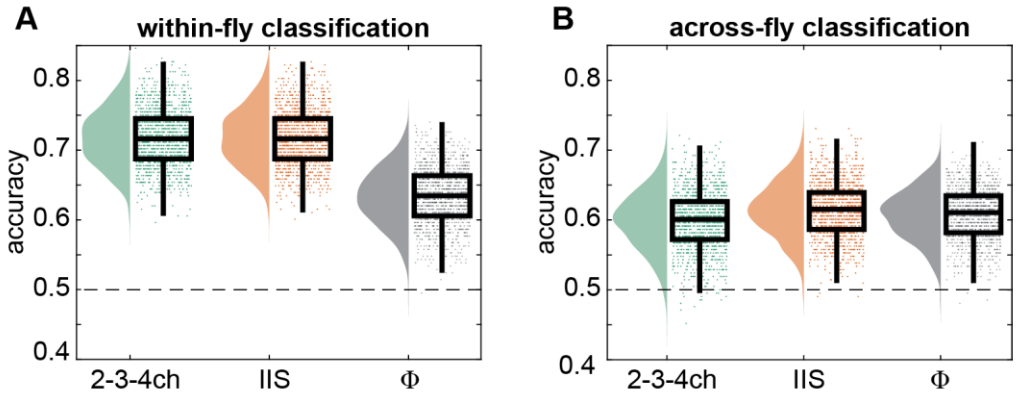


**Fig S7.** Classification accuracy between wakeful vs. anesthetized conditions using the IIS without 1-channel integrated information (green). (**A**) Within-fly and (**B**) across-fly classification. We replot the same results for the IIS and system-level integrated information for comparison (the same data as in Fig 5A-B).

For within-fly classification, the restricted IIS consisting of only 2-, 3-, and 4-channel mechanisms did not achieve significantly different performance to the full IIS ($\chi^2(1) = 0.5691$, $p = 0.451$ using LME model (5), see Methods, where feature had two levels, restricted IIS, i.e. lacking 1-channel mechanisms, or full IIS including all mechanisms; pairwise comparison of restricted IIS to full IIS, $\beta = 9.862 \times 10^{-5}$, $t(7) = -0.658$, $p = .523$; Fig. S7a). For across-fly classification, the restricted IIS achieved worse performance than the full IIS ($\chi^2(1) = 306.5$, $p < .001$; $\beta = -0.0155$, $t(7) = -4.401$, $p = .003$). Taken together with the AIC results (Text S5), we conclude that while 1-channel mechanisms contributed to the IIS, they were not driving its classification performance.
