## Supplementary material for "Integrated information structure collapses with anesthetic loss of conscious arousal in *Drosophila melanogaster*": Text S7

**Text S7. Recurrent connectivity is required for greater system-level integrated information**

In the main text, we infer that recurrent connections throughout the fly brain is reduced by general anesthesia based on our observation that integrated information is reduced during anesthesia. A potential concern is that integrated information may be high in a nonlinear system even in the absence of recurrent connections. Here we provide a simulation to demonstrate that recurrent connectivity is required for greater system-level integrated information.

Here, we compare 2-channel integrated information among 10 simulation runs of 3 auto-regressive models with a nonlinear component: 1) a model with 2 channels that are not physically connected, 2) a model with 2 channels where one channel sends output to the other unidirectionally through a physical connection, and 3) a bidirectionally connected model (the model specifications are given below). Given these models, we would expect system-level integrated information to be greater than zero for model 3 and zero for models 1 and 2, as system-level integrated information requires bidirectional connectivity as explained extensively in [1].

The general form of these models is specified as:

- $X_{t+1} = -0.1X_t + AY_t + e_X$
- $Y_{t+1} = -0.1Y_t + BX_t + e_Y$
- Innovations covariance: diagonal 0.5, off-diagonals 0

(1) In the completely disconnected model:

- $A = 0$
- $B = 0$

(2) In the unidirectionally connected model:

- $A = 0$
- $B = 0.9$ if $X_t >$ threshold; 0 otherwise
    - (i.e., $X$ only influences $Y$ if $X$ is above a certain threshold)
- threshold = 0.9

(3) In the bidirectionally connected model:

- $A = 0.9$ if $Y_t >$ threshold; 0 otherwise
- $B = 0.9$ if $X_t >$ threshold; 0 otherwise
- threshold = 0.9

We compute system-level integrated information on the simulated time series in the same way as in the main text: we 1) binarise the simulated time series based on the median, 2) obtain a TPM, then 3) use PyPhi to compute integrated information, which involves several steps as described in the main text (Fig 1 and [2]).

We find that system-level integrated information is, as expected, much greater for the bidirectionally connected model than the other two models (which are much closer to 0; Fig S8). As integrated information is always above or equal to 0, it is positively biased. While here we included a simple nonlinearity in our model (thresholds), further work should be conducted to assess the behaviour of integrated information also in partially observed systems and non-markovian systems approximated through a Markovian assumption, where spurious high-order correlations might affect the measure.

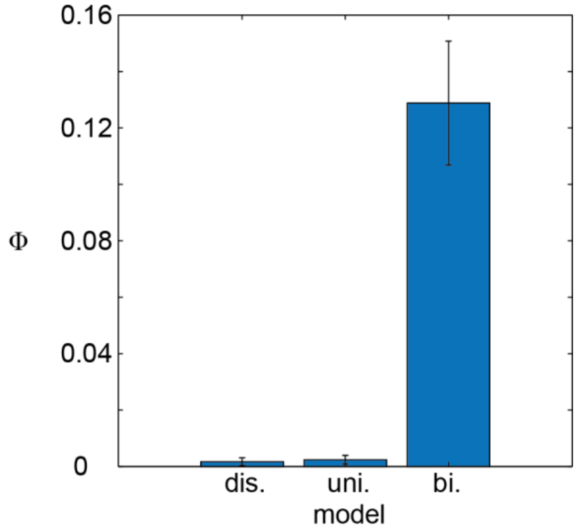


**Fig S8.** System-level integrated information for three simple nonlinear autoregressive models. System-level integrated information is close to 0 when the system is disconnected or unidirectionally connected. Meanwhile, system-level integrated information is much greater than 0 for the bidirectionally connected system. Shown are mean and standard deviation across 10 simulation runs of each model. For each run, a TPM was built such that each row of the TPM was obtained from observing 200 state transitions. We used these TPMs to compute system-level integrated information in the same way as we describe in the main text.
