## Supplementary material for "Integrated information structure collapses with anesthetic loss of conscious arousal in *Drosophila melanogaster*": Text S8

**Text S8. Relation between 1-channel mechanisms and autocorrelation**

In the main text, we consider 1-channel mechanisms to be unclear theoretical constructs of IIT. One possible interpretation is that 1-channel integrated information reflects self-connectivity. Here we quantified the contribution of autocorrelation to 1-channel integrated information.

We directly compared differences (wake minus anesthesia) in 1-channel φ and difference in single-channel autocorrelation (Fig S9). To compute autocorrelation for a given channel, we correlated each LFP time series (of 2.25 s) with itself, shifted by $\tau$ = 4 ms (corresponding to $\tau$ = 4 ms for our integrated information results). Fig S9A plots autocorrelation values against 1-channel φ values for one fly during wakefulness. Note that each channel only has one autocorrelation value but multiple 1-channel φ values (each from a different set of 4 channels; 14 choose 3 = 364 channel sets containing the channel; error bars in Fig S9A are standard deviations across 364 1-channel φ values). Thus, some fixed autocorrelation value (x-axis of Fig S9A) of a given channel corresponds to multiple, highly varied 1-channel φ values (y-axis). This is expected theoretically, because 1-channel φ has to reflect on how the channel is embedded in and interacts with the other three channels.

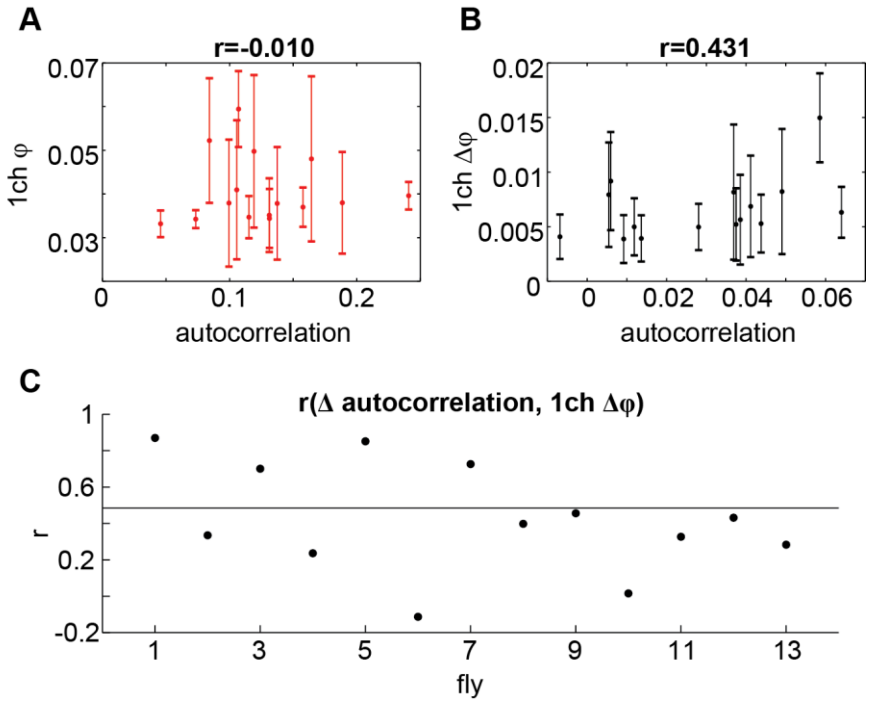


**Fig S9.** Relationship between 1-channel integrated information and autocorrelation, at $\tau$ = 4 ms. (**A**) Single channel autocorrelation plotted against 1-channel integrated information, for a representative fly during wakefulness. Each point corresponds to 1-channel. Error bars are standard deviations of 1-channel φ for a given channel (each channel is contained in 364 out of all 1365 sets of 4 channels). Title gives the correlation coefficient between autocorrelation and 1-channel φ for the fly. (**B**) Difference (wake - anesthesia) in Fisher *z* transformed single-channel autocorrelation (Δ autocorrelation) plotted against difference in 1-channel integrated information (Δφ), for the same fly. Title gives the correlation coefficient between Δ autocorrelation and Δφ for the fly. (**C**) Correlation coefficients between Δ autocorrelation and Δφ for each individual fly. Solid line indicates the average correlation coefficient across flies (coefficients were averaged after Fisher *z* transform, plotted is inverse transform of the mean).

We next subtracted Fisher *z* transformed autocorrelation values during anesthesia from those during wakefulness (Δ autocorrelation). Fig S9B shows Δ autocorrelation plotted against Δφ (wake φ minus anesthetized φ), for the same fly as Fig R3-1A. Correlations at each fly, between Δ autocorrelation and average Δφ values of each channel, indicated that there is some positive correlation between the two measures at the group level (Fig S9C). We confirmed this using a one-sample t-test comparing Fisher *z* transformed correlation coefficients to 0 ($M = 0.424$, $SD = 0.443$, $t(12) = 4.308$, $p = .001$).

In sum, while there seems to be some relationship between the two measures, we conclude that 1-channel integrated information reflects something above and beyond its autocorrelation, namely, the informational and (statistical) causal interactions between that channel with the rest of the channels in the considered system. Whether this is an ideal property for integrated information may need further theoretical exploration in the future.
